# Rethinking Platelet Identity from an Extracellular Vesicle-Centric View: A Proof-of-Concept Study

**DOI:** 10.64898/2026.09.08.750267

**Authors:** Guang Peng, Yan Tian, Yunmin Zhu, Ruoyu Ling, Biao Cheng

## Abstract

The long-standing ambiguity surrounding whether platelets are cells or particles has hindered a unified understanding of their biology. In this study, we characterized the morphology of platelet-like particles (PLPs) using nanoparticle tracking analysis, transmission electron microscopy, and immunofluorescence. Notably, we observed that platelets can be endocytosed by living cells. We also indicate that platelets express the canonical extracellular vesicles (EVs) markers CD9, CD63, CD81, and TSG101. Our results demonstrate that platelets share six key similarities with EVs—in size, morphology, surface markers, function, biogenesis, and mode of action—leading us to propose a novel classification. Our findings suggest that platelets fulfill the current minimal criteria for EVs, raising the possibility that they could be conceptually viewed as a specialized EVs subset rather than conventional blood cells. Adopting this EVs-centric perspective could broaden the current conceptual framework for EVs and offer an alternative lens for interpreting platelet functions, which may in turn help guide future translational research and clinical exploration.

## Introduction

More than a century after Professor Giulio Bizzozero first identified platelets as distinct blood particles, the scientific community has yet to reach a consensus on whether platelets should be classified as living cells[1]. Platelets are anucleate blood components enclosed by a bilayer phospholipid membrane and originate from megakaryocytes. Although often simplistically described as haemocytes—reflecting their well-established role in vascular integrity—platelet functionality extends beyond hemorrhage control and initiation of vascular repair [2, 3]. They actively participate in wound healing and immune regulation through the expression and secretion of cytokines and RNA transcripts. While megakaryocytes and platelets are considered unique to mammals[4], birds and reptiles rely on nucleated thrombocytes for hemostasis [5]. Intriguingly, recent studies have identified anucleate platelet-like fragments in certain invertebrates (e.g., sea star Patiriapectinifera) that facilitate wound repair and immune responses [6]. These fragments resemble extracellular vesicles and express multiple platelet-related markers. The evolutionary emergence of nucleated megakaryocytes and anucleate platelets in mammals may represent an adaptive strategy to maintain vascular integrity more efficiently.

Platelets have long been associated with extracellular vesicles. Historically, extracellular vesicles were termed “platelet dust,” echoing the early description of platelets as “cellular dust” [7]. Indeed, the promotion of coagulation, a critical function of platelets, is also a shared capability of tissue factor-exposing extracellular vesicles that bind factor VII/VIIa[8, 9]. Extracellular vesicles are composed of bilayer phospholipid membranes and comprise diverse subtypes, including exosomes (30–150 nm in diameter), microvesicles (100–1000 nm), apoptotic bodies (100–5000 nm), and oncosomes (1–10 μm)[10–12]. Notably, platelets, with a diameter of 2–4 μm and a thickness of approximately 0.5 μm, exhibit dimensional similarities to apoptotic bodies/oncosomes and microvesicles, respectively [13]. The classification of extracellular vesicles continues to evolve, with novel subtypes continually identified. Recently, a class of giant extracellular vesicles termed blebbisomes has been discovered. With an average diameter of approximately 20 μm—and occasionally exceeding 50 μm—blebbisomes represent the largest extracellular vesicles known to date, surpassing platelets in size[14]. These structures are secreted by various cell types, such as fibroblasts, cardiomyocytes, and cancer cells, through a blebbing process analogous to platelet generation by megakaryocytes. Structurally, blebbisomes share similarities with platelets in their organelle composition, being enriched with components including the Golgi apparatus, endoplasmic reticulum, mitochondria, and ribosomes [15]. Although anucleate, blebbisomes exhibit cell-like characteristics and contain an internal multivesicular endosome (MVE) structure, enabling sustained release of smaller extracellular vesicles—such as exosomes and microvesicles—mirroring the generation of platelet-derived extracellular vesicles (PDEVs) [14].

In fact, platelets share multiple morphological, phenotypic, and functional similarities with extracellular vesicles. Liu et al. reported that circulating tumor cells (CTCs) can internalize entire platelets and acquire the RGS18 gene. High-throughput sequencing further revealed that CTCs contain substantial levels of platelet-derived molecules such as PPBP and PF4[16]. This process resembles the uptake of extracellular vesicles by recipient cells, a key step in realizing EVs-mediated biological functions. Notably, platelets express several evolutionarily conserved extracellular vesicle markers, including CD63 and CD9. CD63, a major platelet activation marker [17], is predominantly localized to lysosomal and dense granule membranes in resting platelets. Upon activation, CD63 translocates to the platelet surface, enhancing activation, spreading, and aggregation[18]. CD9, a key marker expressed in megakaryocytes and platelets, is involved in thrombopoiesis, platelet aggregation, and activation[18, 19]. It forms complexes with integrin αIIbβ3 to modulate platelet adhesion and aggregation[20], contributing to hemostasis and thrombosis. Clinically, impaired CD9 expression or defective platelet aggregation has been associated with diffuse ecchymotic skin lesions and severe thrombocytopenia[21]. Additionally, Annexin A1 and ARF6—recognized markers of microvesicles and oncosomes—are also expressed in platelets [22, 23], where they participate in platelet activation, thrombosis, endocytosis, and platelet-mediated inflammatory responses [24–26]. Therefore, re-conceptualizing platelets as a form of extracellular vesicle may offer an alternative conceptual basis for exploring their non-classical functions and inspire novel therapeutic strategies that harness the bioactive properties of extracellular vesicles.

## Materials and Methods

### Cell culture

MEG-01 and A549 cell lines were purchased from Pricella (CL-0498 and CL-0016, China) and cultured in RPMI 1640 medium (11875093, Gibco,Canada) and Ham’s F-12K medium (21127022,Gibco,Canada) respectively containing 10% (v/v) FBS and 1% penicillin-streptomycin solution(PB180120, pricella, China). Human umbilical vein endothelial cells(HUVECs) were obtained and cultured as described in our previous study[27]. The human dermal fibroblasts (HDF) were obtained from postoperative excess skin tissue of patients undergoing plastic surgery. All experiments have received ethical approval from the Ethics Committee of the general hospital of southern theater command of PLA, and all patients provided written informed consent. HDF were cultured in fibroblast medium (2301, ScienCell, USA). All cell lines were incubated at 37°C in a humidified atmosphere with 5% CO2.

### The isolation of platelet-like particles (PLPs)

MEG-01 cells were treated with 100ng/ml recombinant human thrombopoietin (P00192,Solarbio,China) and PMA (P6741,Solarbio,China) for 5 days as reported [28]. The cell culture supernatant was collected and centrifuged at 800g for 15 minutes. The pellet were discarded and the supernatant was again centrifuged at 1600g for 15 minutes, then the precipitation containing PLPs was suspended with PBS. The concentration of PLPs was detected by a hematology analyzer (Mindray, China).

### The isolation of platelet

Peripheral blood samples were collected from healthy donors and utilized to obtain platelet as described in our previous research [12]. In brief, fresh peripheral blood was collected in 8.5ml tube contain ACD-A (364606, BD, USA) and centrifuged at 200g for 15 minutes. Platelet-rich plasma (PRP) was isolated and transferred into tube containing 1/10(v/v) ACD-A and 100 ng/ml PGE1(745-65-3,Sigma-Aldrich,USA).PRP was again centrifuged at 900g for 15 minutes and supernatant was aspirated. Following two washes with Hepes-NaCl buffer, the platelet pellet was resuspended in Tyrode-Hepes buffer. The platelets concentration were measured by a hematology analyzer (Mindray, China). All experiments received ethical approval from the Ethics Committee of the general hospital of southern theater command of PLA.

### Nanoparticle tracking analysis (NTA)

The number, size distribution of PLPs was measured by Nanosight NS300(Malvern panalytical,UK) as described in our previous study [12].

### Transmission electron microscope (TEM)

An appropriate amount of PLPs or platelet suspension was applied onto a TEM copper grid and allowed to adsorb at room temperature for 5 min. Excess liquid was carefully removed using filter paper. Uranyl acetate solution was then dropped onto the grid to cover the sample for negative staining, which lasted for 5 min. After removing the staining solution with filter paper, the grid was air-dried at room temperature. Images were acquired using an HT-7700 transmission electron microscope operated at 100 kV and subsequently analyzed.

### Immunofluorescence analysis

Immunofluorescence analysis was performed as described in our previous research [12]. In short, HDF, HUVECs and A549 cells were fixed with paraformaldehyde (4%) and the cell membranes were permeabilized via Triton X-100(0.3%). Goat serum was added to blocking prior to the staining procedure. The cells were incubated overnight at 4°C with antibodies against CD41(ab134131, abcam, UK). Slides were washed in PBS three times and then incubated for 1 hour using donkey anti-rabbit IgG H&L (Alexa Fluor® 488) secondary antibody (ab150073, abcam,UK) at room temperature(RT).The cells were stained with CoraLite® Plus 488 Phalloidin(PF00001,proteintech,China) for 30min at RT to visualize filamentous actin. Nuclei were visualized by DAPI staining (ab104139, abcam,UK). Images were captured using a fluorescence microscope.

### PLPs/PLT in vitro uptake experiment

For PLPs/PLT in vitro uptake experiments, 1/20(v/v)10^12^/ml of DiD-labeled PLPs/PLT was cocultured with HDF, HUVECs or A549 cells and incubated for 24 h. Following three washes with PBS buffer, the endocytosis of DiD-labeled PLPs/PLT was observed by immunofluorescence analysis. Images were captured using Nikon AX/AX R confocal scanning laser microscope (Nikon,Japan).

### Western blot analysis

Western blot analyses were performed as described in our previous research [29]. The following antibodies were used: CD9(F1178, selleck), TSG101(F1543, selleck), CD63(ab134045, abcam), CD81(R381296, zenbio), CD41(HA722940, huabio).

### Optical Diffraction Tomography (ODT) dual-modality imaging

Super-resolution imaging of nucleolar structures was performed using a commercial microscope (Cell Xtreme, CSR Biotech). Images were acquired using a 100×/1.5 NA oil immersion objective (Evident). HUVEC cells were seeded in 8-well chambered coverglass and maintained at 37℃ and 5% CO2 in a humidified chamber for live ODT imaging. Then adherent HUVEC cells were cocultured with platelets labeled with DID(Green).ODT images were collected and analyzed as described previously [30],and sparse deconvolution was carried out to further improve the image quality [31].

## Results

Given the close relationship between the origin and function of extracellular vesicles and their parent cells, along with the current difficulty in fully replicating megakaryocyte-driven platelet production in vitro, we focused on megakaryocyte-derived extracellular particles to identify evidence supporting the classification of platelets as extracellular vesicles. Herein, we generated platelet-like particles (PLPs) from the human megakaryocytic cell line MEG-01 using thrombopoietin (TPO) and phorbol 12-myristate 13-acetate (PMA), as previously described, to model platelet-like functions[28]. During PLP production, the majority of suspended MEG-01 cells differentiated into adherent megakaryocytic cells exhibiting long, branching membrane projections and a marked increase in cell size (**Figure 1A**). Since alterations in microstructure often reflect functional changes, we next sought to characterize the morphology of the generated PLPs.We employed nanoparticle tracking analysis (NTA) and transmission electron microscopy (TEM) to assess the size distribution and ultrastructure of the PLPs, respectively (**Figure 1B-C**). The PLPs appeared as membrane-bound vesicles with a typical bilayer structure and a median diameter of 183 nm, which falls within the size range of exosomes. Notably, the centrifugal force required to pellet PLPs was only 1,600 ×g, whereas exosome isolation in our previous work required forces exceeding 100,000 ×g[12]. As particle internalization by recipient cells is essential for the functional delivery of EVs cargo, we incubated DiD-labeled PLPs with several human cell lines and subsequently stained them with the platelet-specific marker CD41. As expected, PLPs were efficiently taken up by human primary fibroblasts, vascular endothelial cells, and cancer cells (**Figure 1D**), suggesting the understudied impact of PLPs on specific cell types. Conversely, the inclusion of dynasore, a potent endocytosis inhibitor, diminished the uptake process of PLPs by target cells (**Figure 1E**).

**Figure 1.**
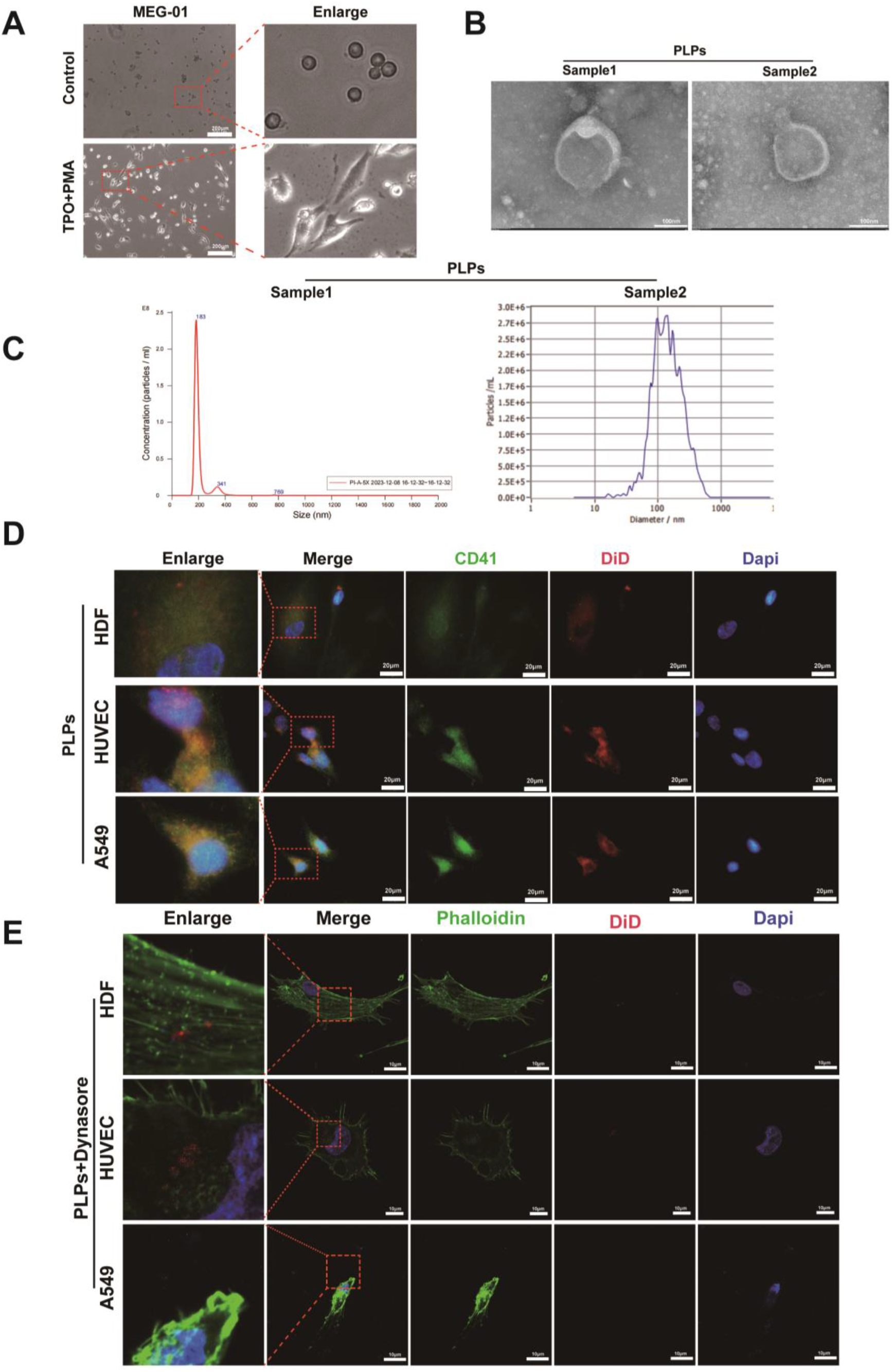
**A**. Morphological changes of MEG-01 treated with TPO and PMA. Scale bar =200μm. **B-C**. Size distribution of PLPs detected by transmission electron microscopy (B, scale bar =100nm) and NTA (C) respectively. **D**. Immunofluorescence staining micrographs of CD41 in human derived fibroblast (HDF), HUVEC and A549 cocultured with DID labeled platelet-like particles (PLPs). Scale bar =20μm. **E**. Representative immunofluorescence staining micrographs of dynasore treated human derived fibroblast (HDF), HUVEC and A549 cocultured with DID labeled platelet-like particles (PLPs). Scale bar =10μm.

Although PLPs are closer in size to exosomes, their budding pattern and CD41 expression in MEG-01 cells establish them as a simplified model for investigating megakaryocyte-derived extracellular vesicle/platelet-like particle biogenesis. We acknowledge that PLPs do not fully recapitulate platelets; nevertheless, they reinforce the concept that megakaryocyte-shed particles exhibit EVs properties. However, given that the average size of PLPs is substantially smaller than that of intact platelets (2–4 μm), we conducted subsequent experiments using platelets rather than PLPs. Firstly, we conducted a detailed morphological analysis of donor-derived platelets using immunofluorescence and transmission electron microscopy (TEM) to measure their diameter and evaluate structural features (**Figure 2A-B**). As surface marker expression constitutes a key criterion for EVs identification, we assessed multiple classical EV markers in activated and resting platelets using Western blot (**Figure 2C**). Consistent with their dual identity, platelets expressed the common markers CD9 and CD63. Intriguingly, they also tested positive for CD81 and TSG101, markers often regarded as more specific to EVs. Since the endocytic uptake of extracellular vesicles (EVs) by recipient cells is critical for content delivery and functional modulation, we further evaluated the internalization of platelets by various cell types. Notably, using confocal microscopy, we captured a dynamic process of recipient cells internalizing DiD-labeled platelets (**Figure 2D**). Furthermore, CD41-labeled platelets were efficiently taken up by human dermal fibroblasts (HDFs), human umbilical vein endothelial cells (HUVECs), and A549 cells (**Figure 2E**). The dual-modality Optical Diffraction Tomography (ODT) imaging enables intuitive visualization of the endocytosis of DID-labeled platelets by HUVECs (**Figure 2F, Supplementary Video 1**).

**Figure 2.**
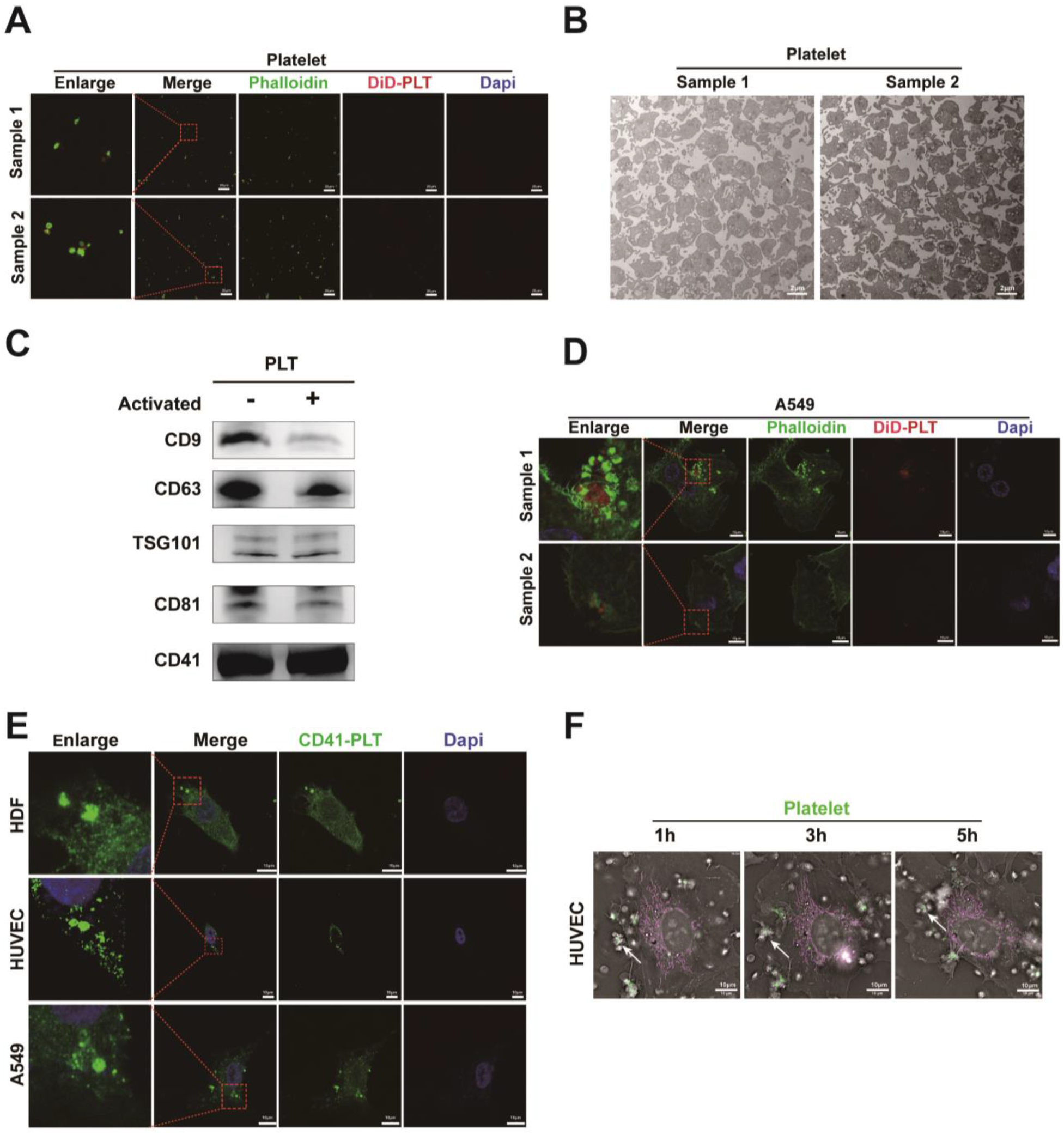
**A**. Representative immunofluorescence staining micrographs of platelet stained by DID and phalloidin. Scale bar =10μm. **B**. Representative transmission electron microscopy images of platelet. Scale bar =2μm. **C**. Western blot analysis was performed to assess the expression of classical extracellular vesicle markers (CD9, CD63, CD81, TSG101) and the platelet marker CD41 in activated and unactivated platelets. **D**.The process of A549 cells endocytose platelet stained by DID, actin filaments was labeled by phalloidin. Scale bar =10μm. **E**. Representative immunofluorescence staining micrographs of CD41 in human derived fibroblast (HDF), HUVEC and A549 cocultured with platelet. **F**. Live-cell imaging of HUVEC cells endocytosing platelets labeled with DID(Green). Mitochondria were labeled with mitochondrial fluorescent dye (PKMITO^®^ Red). Scale bar =10μm.

We also confirmed that DiD-labeled platelets were internalized by HDFs, HUVECs, and A549 cells (**Figure 3A**). However, treatment with the endocytosis inhibitor dynasore markedly reduced platelet uptake (**Figure 3B**), supporting an endocytosis-dependent mechanism for this process. Collectively, these findings suggest that platelets can exert EVs-like functions across a range of physiological contexts.

**Figure 3.**
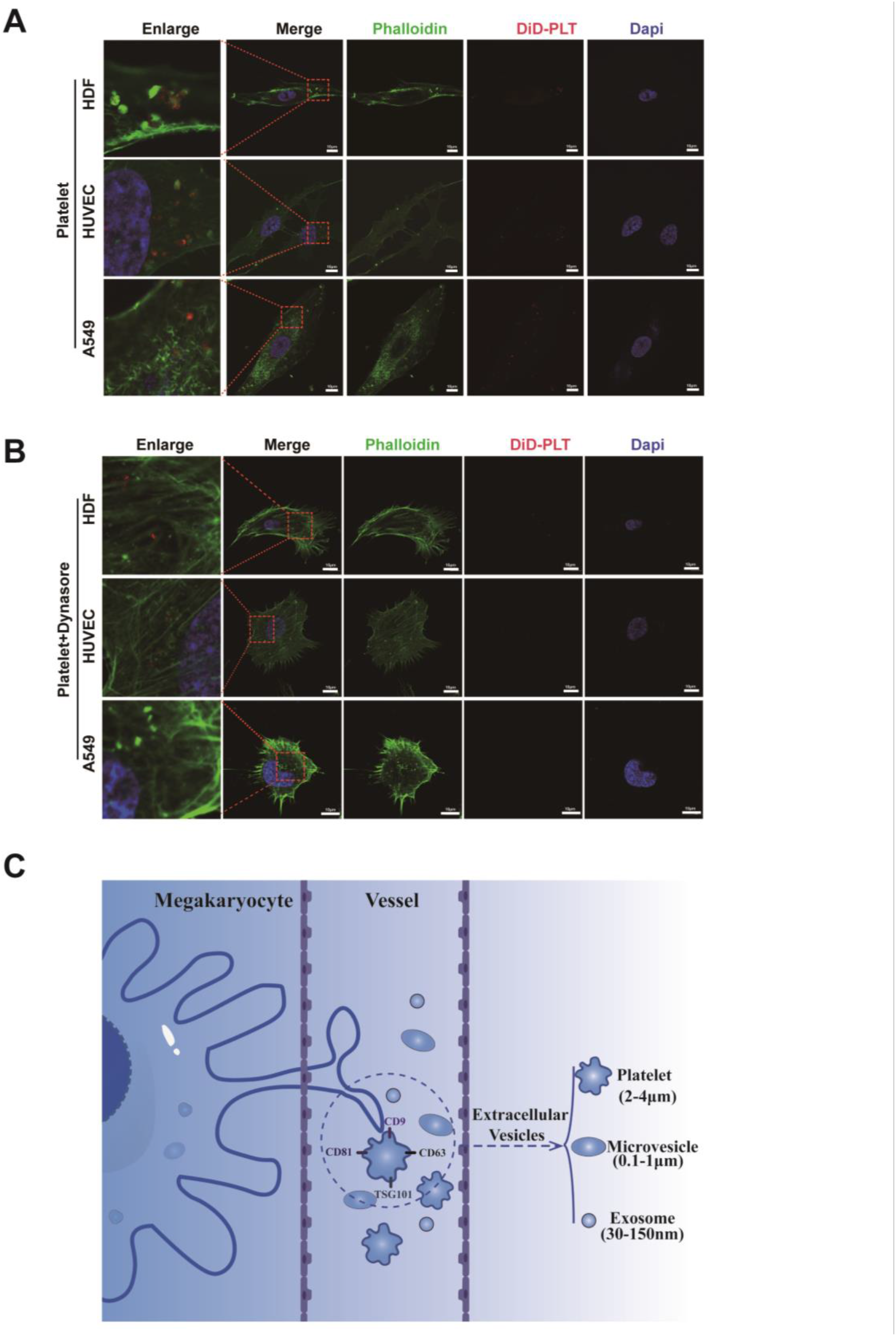
**A**. Immunofluorescence staining images of fibroblast (HDF), HUVEC and A549 cells cocultured with DID labeled platelet. Scale bar =10μm. **B**. Immunofluorescence staining images of dynasore (15μm) treated fibroblast (HDF), HUVEC and A549 cells cocultured with DID labeled platelet. Scale bar =10μm.**C**. Scheme for the production of platelet, microvesicle, exosomes in the bone marrow microenvironment. Platelets can be categorized as megakaryocyte-derived EVs (2-4μm) with diverse physiological functions, expressing the canonical EVs markers CD9, CD63, CD81, and TSG101.

## Discussion

A refined classification of heterogeneous platelets would benefit from a prior understanding of megakaryocyte heterogeneity and their mechanisms of platelet production. Megakaryocytes can be broadly classified into platelet-producing, niche-supporting, and immune-regulatory subtypes[32]. EVs secretion represents a key pathway through which megakaryocytes perform immunomodulatory and niche-supporting roles [33, 34]. Notably, the process of platelet biogenesis shares several similarities with EVs release. The mechanism of platelet formation has been debated for decades. Using whole-organ 3D and 4D quantitative imaging, Potts et al. reported that megakaryocytes release proplatelets via membrane budding, suggesting this as the major mode of platelet biogenesis[35]. However, while budding megakaryocytes account for 70%–80% of those in bone marrow, proplatelets are detected at a considerably lower frequency of merely 2% [36]. Italiano et al. characterized these membrane buds as cell fragments and highlighted the difficulty in morphologically distinguishing megakaryocyte-derived extracellular vesicles (MKEVs) from proplatelets, given their structural and functional similarities[37]. In contrast, Jackson and colleagues proposed that megakaryocyte buds are distinct from EVs and may represent intermediate stages in platelet formation [36]. Further mechanistic insight was provided by Ellis et al., who demonstrated that the size and spatial distribution of megakaryocyte buds—including developing platelets—are regulated by interactions between glycoprotein Ibα (GPIbα) and filamin A (FlnA)[38], which may be essential for their maturation into functional platelets. Beyond structural resemblances, platelets and EVs also exhibit functional overlap. Certain MKEVs express elevated levels of platelet markers such as CD41, phosphatidylserine, and CD62P [39], which contribute to thrombus formation and megakaryopoiesis [37]. These findings suggest that proplatelets/platelets and MKEVs may share a common origin, though the precise relationship remains unclear. Naturally, further in-depth research is warranted to clarify how to identify and distinguish conventional extracellular vesicles from megakaryocyte-derived vesicles that exhibit platelet-like functions.

In summary, when assessed against the minimal criteria for extracellular vesicles established by the International Society for Extracellular Vesicles (ISEV) [40], platelets exhibit a set of defining characteristics that support their classification as EVs:1. Platelets are anucleate and delimited by a phospholipid bilayer [3];2. Platelets are secreted from living cells, lacking the capacity for self-replication [3];3. Their size distribution (2-4μm) falls within the range of many conventional EVs populations;4. Platelets express typical EVs-associated surface markers, including CD9, CD63, CD81, TSG101 [17–20]; 5. Functionally, platelets can be internalized by recipient cells, thereby transferring bioactive molecules, or can modulate cellular activities via specific ligand–receptor interactions[41];6. Beyond hemostasis, platelets act as active communicators that regulate immune and inflammatory responses, analogous to EVs[42]. With aging, their function transitions toward a proinflammatory and immunomodulatory phenotype [43]. Collectively, these attributes provide a strong rationale for considering platelets as a naturally produced, multifunctional subtype of extracellular vesicles. Herein, we demonstrate that platelet-like particles exhibit key EVs-like morphological characteristics, offering a valuable model to study platelet physiology and their functional resemblance to extracellular vesicles. We further demonstrate that platelets can be internalized by various human cells via endocytosis, analogous to EVs uptake. Moreover, we identify that platelets express the canonical EVs markers (CD9, CD63, CD81, TSG101), suggesting an EVs-like role in intercellular communication.

Conversely, platelets also exhibit distinct characteristics that set them apart from extracellular vesicles. These include the ability to perform limited, non-sustained synthesis of specific proteins using megakaryocyte-derived mRNA[44], as well as the capacity to shift their energy metabolism from glycolysis to mitochondrial oxidative phosphorylation upon activation, thereby enabling rapid energy supply. Such metabolic flexibility allows platelets to respond quickly at sites of vascular injury and fulfill their hemostatic function[45]—features not observed in conventional extracellular vesicle. Although platelets display phenotypes and characteristics distinct from both intact cells and extracellular vesicles, they appear to satisfy only the current minimum criteria for EVs rather than those defining cells. Based on these findings, we propose the reclassification of platelets as a unique category of EVs produced by megakaryocytes, analogous to other megakaryocyte-derived exosomes and microvesicles (**Figure 3C**).

We acknowledge several limitations. First, the PLPs generated from MEG-01 cells are considerably smaller than native platelets, rendering them an imperfect model for recapitulating authentic platelet biogenesis but a useful surrogate for studying EVs-like budding. Second, our uptake experiments were exclusively performed in vitro; whether such endocytic events occur in vivo under hemodynamic flow and what downstream functional consequences arise in recipient cells warrant further investigation. Third, while we detected EVs markers (CD9, CD63, CD81, TSG101) in platelet lysates, we did not distinguish their surface versus intracellular pools via flow cytometry or immunogold labeling. Given that CD63 resides primarily in granule membranes in resting platelets, its classification as a “platelet EVs surface marker” requires cautious interpretation. Such comparative studies are essential to firmly establish functional equivalence prior to any reclassification. Despite these caveats, our findings strongly support the notion that platelets satisfy the current minimal criteria for EVs and offer a new framework for rethinking platelet biology from an EVs-centric perspective. Therefore, viewing platelets from the perspective of extracellular vesicles will not only help elucidate their origin, function, and relationship with megakaryocytes but also inform improvements in platelet isolation and preservation protocols, the exploration of novel applications, and the identification of new clinical indications.

## Supporting information

visualization of the endocytosis of DID-labeled platelets by HUVECs (Figure 2F, Supplementary Video 1)

## Abbreviations

EVs: extracellular vesicles
PLPs: platelet-like particles
PLT: platelets
MVE: multivesicular endosome
PDEVs: platelet-derived extracellular vesicles
CTCs: circulating tumor cells
NTA: Nanoparticle tracking analysis
TEM: Transmission electron microscope
ODT: Optical Diffraction Tomography
TPO: thrombopoietin
PMA: phorbol 12-myristate 13-acetate
HDFs: human dermal fibroblasts
HUVECs: human umbilical vein endothelial cells
MKEVs: megakaryocyte-derived extracellular vesicles

## Ethics Approval

The study was conducted in accordance with the Declaration of Helsinki, and has received ethical approval from the Ethics Committee of the general hospital of southern theater command of PLA.

## Author Contributions

Guang Peng conceptualized the study, designed and performed the experiments, analyzed the data and wrote the manuscript. Biao Cheng conceptualized and supervised the study, designed the experiments, and critically revised the manuscript. Yan Tian,Yunmin Zhu and Ruoyu Ling performed part of experiments and revised the manuscript.

